# Hidden paths to endless forms most wonderful: Parasite-blind diversification of host quality

**DOI:** 10.1101/2020.12.23.424158

**Authors:** Lisa Freund, Marie Vasse, Gregory J. Velicer

## Abstract

Evolutionary diversification can occur in allopatry or sympatry, can be unselected or driven by selection, and can be phenotypically manifested immediately or remain phenotypically latent until later manifestation in a newly encountered environment. Diversification of host-parasite interactions is frequently studied in the context of intrinsically selective coevolution, but the potential for host-parasite interaction phenotypes to diversify latently during parasite-blind evolution is rarely considered. Here we use a social bacterium experimentally adapted to several environments in the absence of phage to analyse allopatric diversification of latent host quality - the degree to which a host population supports a viral epidemic. Phage-blind evolution reduced host quality overall, with some bacteria becoming completely resistant to growth suppression by phage. Selective-environment differences generated only mild divergence in host-quality. However, selective environments nonetheless played a major role in shaping evolution by determining the degree of stochastic diversification among replicate populations within treatments. Ancestral motility genotype was also found to strongly shape patterns of latent host-quality evolution and diversification. These outcomes show that adaptive landscapes can differ in how they constrain stochastic diversification of a latent phenotype and that major effects of selection on biological diversification can be missed by focusing on trait means. Collectively, our findings suggest that latent-phenotype evolution (LPE) should inform host-parasite evolution theory and that diversification should be conceived broadly to include latent phenotypes.

---

Wonderful “endless forms” of phenotypes^1–3^ often first evolve non-adaptively^4–7^, even if they later prove beneficial in a new context as “exaptations”^8^. Temporally, a non-adaptively evolved phenotype might be generated immediately when its causal genotype first evolves, or only later, upon that genotype’s exposure to a novel or changed environment. Here we refer to a phenotype that is potentiated by an existing genotype but not initially manifested due to environmental specificity as a ‘latent phenotype’. We additionally refer to the evolution of such a genotype prior to manifestation of its initially latent phenotype as ‘latent-phenotype evolution’ (or ‘LPE’), a label independent of the causes or consequences of LPE (see Methods).

LPE is intrinsically non-adaptive because the focal phenotype is not yet generated when its genetic basis first evolves. However, the genetic basis of the latent phenotype might evolve by any evolutionary mechanism (Table 1). For example, the causal genotype might arise adaptively due to one phenotypic effect beneficial in a first environment while pleiotropically potentiating a second phenotypic effect that is only manifested in a distinct environment encountered later^9–10^. Alternatively, genotypes underlying LPE may evolve non-adaptively by hitchhiking with an adaptive genetic element^11^ or by stochastic forces^6,12^. Variation for initially neutral alleles underlying latent phenotypes^12,13^ has long been recognized as potential fuel for later adaptation to new and changing environments^14–16^.

**Table 1.** Categories of latent-phenotype evolution (LPE). The genetic basis of a focal latent phenotype first evolves in a temporally prior environment E_*Pr*_. We distinguish three mechanisms potentially responsible for evolution of the causal genotype: adaptation, hitchhiking and stochasticity (corresponding to alleles *a, h* and *s*, respectively). By definition, each allele potentiates a phenotype *y* that is latent in the prior environment E_*Pr*_ (latent phenotype *y, LP_y_*, right superscript) but is then manifested in the later-encountered environment E_*L*_ (manifested phenotype *y*, MP_*y*_, right superscript). In addition, this allele may cause a non-advantageous manifested phenotype *x* in the prior environment E_*Pr*_ (MP_*x*_, left superscript), that might or might not also be manifested in E_*L*_ (signified by parentheses). All phenotypes manifested in E_*L*_ can have a positive, a negative, or no fitness effect. The first (non-header) row of the table describes a form of pleiotropy - a delayed, environment-contingent pleiotropic effect of an adaptive allele^61^. The rows below describe scenarios consistent with the common meaning of ‘cryptic genetic variation’^12^ (see Methods).

| Prior environment $E_{Pr}$ | Later environment $E_L$ | Cause of allele's increase in $E_{Pr}$ | Allele's adaptive status in $E_{Pr}$ |
| --- | --- | --- | --- |
| $MP_x a^{LP_y}$ | $(MP_x) a^{MP_y}$ | adaptation (direct selection) | adaptive |
| $h^{LP_y}$ | $h^{MP_y}$ | hitchhiking (indirect selection) | non-adaptive |
| $MP_x h^{LP_y}$ | $(MP_x) h^{MP_y}$ | hitchhiking (indirect selection) | non-adaptive |
| $s^{LP_y}$ | $s^{MP_y}$ | stochasticity (mutation/drift) | non-adaptive |
| $MP_x s^{LP_y}$ | $(MP_x) s^{MP_y}$ | stochasticity (mutation/drift) | non-adaptive |

Latently evolved phenotypes can be features of individual organisms. For example, bacteria have latently evolved altered antibiotic resistance^10,17^, metabolic-profile shifts^15^ and changes in nutrient-uptake abilities^18^. However, outcomes of between-organism interactions can also be considered phenotypes. Examples of such “interaction phenotypes”^19,20^ include reproductive incompatibility resulting from allopatric speciation, which remains latent until allopatrically diverged lineages make secondary contact^21–23^. Similarly, host-parasite interaction phenotypes^24–28^ and bacterial social-interaction phenotypes^29,30^ can also evolve and diversify latently in allopatry. Given that i) pleiotropy, hitchhiking and genetic drift are common, ii) manifestation of phenotypes is often context-specific (e.g. due to phenotypic plasticity^31^ or limitations on interaction opportunities for interaction phenotypes), iii) exposure to changing or new environments is inevitable for most biological lineages, and iv) latently evolved phenotypes often have selective significance upon their manifestation, LPE is likely to contribute substantially to long-term patterns of phenotypic evolution and diversification.

Bacteria engage in a vast array of interactions, including with their own viral parasites, bacteriophages. Bacteria-phage interactions are determined by the match between phage-infectivity and bacterial-resistance mechanisms, which can result in narrow to broad host ranges^32–34^. Diverse mechanisms to resist phage have evolved at all major stages of the infection cycle: from preventing phage adsorption and impeding post-entry reproduction and assembly to stopping virion release through abortive systems that kill both phage and host^32,35^. Selection to resist infection can lead to host-phage incompatibility, as antagonistic coevolution between phage and their hosts leads to rapid local adaptation^36,37^ and diversification^27,38,39^. However, how much host quality - the degree to which a host genotype or population facilitates parasite growth - is shaped directly by parasite-imposed selection versus indirectly from byproducts of other selective forces (e.g. resource competition^26^ or other predators^40^) or stochastic forces has been little investigated.

Perhaps all microbes express some social traits^41^, but some have evolved extraordinarily complex suites of cooperative behaviours^42^. Myxobacteria, including the model species *Myxococcus xanthus*, engage in cooperative swarming^43^ during group predation^44^ and multicellular fruiting-body development^45^. As predators of many microbes, myxobacteria are predicted to shape the structure and evolution of soil microbial communities^46,47^.

Myxobacteria are themselves subject to selective pressure by myxophage^48^, which in turn are likely to shape myxobacterial social evolution. For example, cell-surface molecules such as type-IV pili and O-antigen serve as phage receptors in many bacteria^49,50^ and also function in *M. xanthus* social behaviours^51,52^. Thus, just as bacteria-phage coevolution can indirectly shape bacterial social interactions^53–55^, social evolution in the absence of phage is likely to latently alter the character and diversity of future host-parasite interactions. Analyses of experimentally-evolved lineages^29,30^ suggest that intra-specific social interactions between natural *M. xanthus* lineages^56^ often evolve latently. But how LPE shapes future antagonistic interactions of myxobacteria with other species, including with phage and their own prey, remains largely unexplored.

In this study, we test for LPE - including diversification - of host-virus interactions using the virulent myxophage Mx1^57^ and bacterial populations from an evolution experiment recently named MyxoEE-3^7^ (see Methods). In MyxoEE-3, populations of *M. xanthus* underwent selection for increasing fitness at the leading edge of colonies expanding spatially by growth and motility, with cells near the leading edge transferred to a fresh plate at the end of each evolutionary cycle^29,30,58^. Importantly, evolving populations never encountered phage during MyxoEE-3, thereby allowing us to test whether and how phage-blind adaptation to multiple environments indirectly shapes the character of interactions with obligate parasites.

We first analyse host-phage LPE in eight MyxoEE-3 treatments that share a common ancestor but differed in selective environment with respect to surface structure (hard vs. soft agar), nutrient availability (high or low nutrient levels) and/or nutrient source (nutrient medium alone or with prey lawns). We examine effects of environment and chance on the direction and degree of average phage-blind host-quality evolution and also on the degree of within-treatment diversification. If LPE is mediated predominantly by alleles that increased due to selection during MyxoEE-3, adaptive landscape structure^59,60^ may shape LPE outcomes, including the degree to which latent phenotypes diversify stochastically.

We further analyse how ancestral motility genotype shapes LPE by testing for effects of each of the two *M. xanthus* motility systems on latent host-quality evolution and diversification.

## Methods

### Semantics and nomenclature

#### On use of ‘latent-phenotype evolution’

Large bodies of literature examine modes by which the genetic basis of latent phenotypes evolves, for example as initially adaptive alleles that potentiate initially unrealized pleiotropic phenotypes^9–61–62^ or as initially neutral alleles, variation for which is commonly referred to “cryptic genetic variation” (CGV)^12,13^. However, there does not appear to be a well-established generic label for the evolution of the genetic basis of latent phenotypes that is impartial with regard to causes or consequences of such evolution and that applies equally to the phenotypes of individuals and between-organism-interaction phenotypes.

We adopt ‘latent’^63^ over ‘cryptic’ due to the frequent association of the latter with selectively neutral alleles^12,13,16^ (despite exceptional applications to initially adaptive alleles^64^) and the evocation of future manifestation by ‘latent’. We use ‘latent-phenotype evolution (LPE)’ to focus primary attention on the process of evolution over time, which may result in loss of variation due to fixation of an allele underlying a latent phenotype, rather than primarily on within-population variation at loci encoding such alleles, which is the focus of CGV. We also note that LPE in our sense is distinct from (but may nonetheless be related to) the latent evolvability of a genotype, population or species, *i.e*. the potential for future evolution of novel forms, functions or diversity^65–68^.

#### On use of ‘manifestation’

Here we use ‘manifestation’ (and variations) rather than ‘expression’ to refer to the actualization of a genetically caused phenotype. This is because we conceive expression to be actualization of a phenotype by an individual organism, but we desire a term that also applies generically to between-organism-interaction phenotypes. Actualization of the latter may be prevented simply by lack of spatial proximity between the relevant organisms rather than by lack of phenotypic expression by individuals.

#### MyxoEE-3

To facilitate reference to the broader evolution experiment of which the treatments examined were a part, we refer to the overall experiment as MyxoEE-3^7^ (Myxobacteria Evolution Experiment, with ‘3’ indicating the temporal rank of the first publication from MyxoEE-3^69^ relative to first publications from other MyxoEEs^7^). Shared features of MyxoEE-3 treatments have been described previously^29,30,58^ Treatments examined here are summarized in Table S1.

### Strains and procedures

#### Strains

In MyxoEE-3, ancestral strains differing in motility genotype and antibiotic-resistance marker were selected for increased fitness at the leading edge of expanding colonies. Multiple treatment-sets of replicate populations adapted to different environmental conditions that varied in nutrient level, nutrient type or agar concentration (Table S1).

The ancestral wild type strains GJV1 and GJV2 have two functional motility systems: adventurous (A) and social (S) motility (hereafter referred to as the A+S+ motility genotype)^70^. Deletion of one gene essential for either motility system led to strains that were defective in A-motility (A-S+; deletion of *cglB* in GJV3 and GJV5) or S-motility (A+S-; deletion of *pilA* GJV4 and GJV6). GJV1, GJV3 and GJV4 are rifampicin-sensitive, whereas GJV2, GJV5 and GJV6 are rifampicin-resistant variants of the corresponding motility type (Table S1). Two distinct sub-clone genotypes of GJV1 (represented by GJV1.1 and GJV1.2) previously found to differ by one mutation were used to establish the rifampicin-sensitive A+S+ MyxoEE-3 populations^29^ and thus were examined here also. No phenotypic differences between these clones were found in our assays (see *Statistical analysis*). A single sub-clone was used for each of the other five ancestral strains since there are no known mutational differences between the ancestral sub-clones used to found the respective MyxoEE-3 populations.

For this study, we used MyxoEE-3 populations that evolved for 40 two-week cycles under high-nutrient conditions (CTT growth medium), low-nutrient conditions (0.1% Casitone-CTT) or with prey (*Escherichia coli* or *Bacillus subtilis* and CTT). Additionally, the agar concentration in each environment was either high (1.5% hard agar, HA) or low (0.5% soft agar, SA) (Table S1). During evolution, replicate populations derived from each of the six ancestors (GJV1-GJV6) grew and swarmed (to their ability) outwards on the surface of each selection environment for two weeks, after which a patch of defined size was collected from the leading edge of each colony and transferred to a fresh plate. Importantly, these populations never interacted with phage during MyxoEE-3.

The virulent myxophage Mx1^57^ is a double-stranded DNA Myoviridae, morphologically similar to coliphages T2 and T4^48,71^. Our stock of Mx1 was generated through infecting a growing liquid culture of GJV1. We isolated phage particles with 10% chloroform and filtration (0.22 μm) and titered the resulting solution using plaque assays. *M. xanthus* strain DZ1^72^ was used for phage-quantification assays.

#### Growth conditions and phage infection

All experiments were performed in liquid CTT medium (10 mM Tris pH 8.0, 8 mM MgSO_2_, 10g/l casitone, 1 mM KPO_4_, pH 7.6) supplemented with 0.5 mM CaCl_2_ in case of phage infection and incubated at 32 °C and 300 rpm. Prior to each experiment, bacteria were inoculated onto CTT 1.5% agar from frozen stocks and incubated at 32 °C and 90% rH until sufficiently grown. Colony-edge samples were transferred to 8 ml CTT liquid and incubated shaken at 300 rpm. When cultures reached mid-log phase, cells were centrifuged (15 min, 5000 rpm) and resuspended with CTT liquid to ~2×10^8^ cells/ml. Phage particles were added to bacterial populations from the same phage stock at ~2×10^6^ particles/ml, resulting in a multiplicity of infection (MOI) of ~0.01. Phages were allowed to infect bacteria for 24h, upon which phage were isolated. To do so, 100 *μ*l chloroform were added to 1 ml of culture and this mixture was incubated for 5 min under constant shaking/vortexing to disrupt bacterial cells. Subsequently, dead cells and phage were separated by centrifugation (3 min, 12000 rpm). Supernatant containing the phage was stored at 4 °C. To quantify the phage number for each bacterial population after 24 h of infection, phage solutions were diluted, 10 μl were mixed with 10 μl indicator strain DZ1 (10^8^ cells/ml) and 1 ml CTT 0.5% agar. This mixture was poured onto 5 ml CTT 1.5% hard agar, plates were closed immediately and incubated until plaque-forming units were visible. We performed four replicates of this experiment, each divided into three randomized blocks.

#### M. xanthus growth in the presence of phage

To assess effects of phage on growth of populations P65-P72 (Table S1), we measured optical density (OD 595 nm) of liquid cultures growing with and without phage. Overnight cultures of each ancestor and evolved population were diluted in equal volumes into two 15 ml cultures in 100 ml flasks, one of which was infected with phage (MOI of 0.01), and incubated shaken as described above. Measurements were taken at 0, 14, 16, 18, 20, 22 and 24 h.

#### Statistical analysis

Details of statistical analysis can be found in the supplementary materials.

## Results

### Phage-blind evolution lowered host quality overall while environment mildly shaped treatment means

Host quality depends on a combination of host features, including extracellular components phage must bypass or penetrate to reach the cell surface^73^, surface and membrane components that phage use for invasion^49^, harmful intracellular components phage must avoid or neutralize^74^, and beneficial intracellular components that phage exploit for growth. Any of these components might be altered during adaptation in the absence of phage. If selective pressures imposed by distinct MyxoEE-3 environments differ in their effects on traits important to phage invasion and growth, the MyxoEE-3 treatments might often vary in host quality. If not, more variation in host quality should be found among replicate populations within treatments than between treatment means.

Evolved populations and their ancestors were exposed to phage epidemics in shaken liquid culture for 24 h before phage population sizes were determined. On average across all 72 populations descended from the A+S+ ancestor, evolved populations supported less phage growth than their ancestors (Fig. 1a; t_54_ = −4.78, *p* < 0.001). Indeed, 28 populations latently evolved to become significantly lower-quality hosts (from the phage perspective) than their ancestors, whereas none supported significantly greater phage growth.

**Figure 1.**
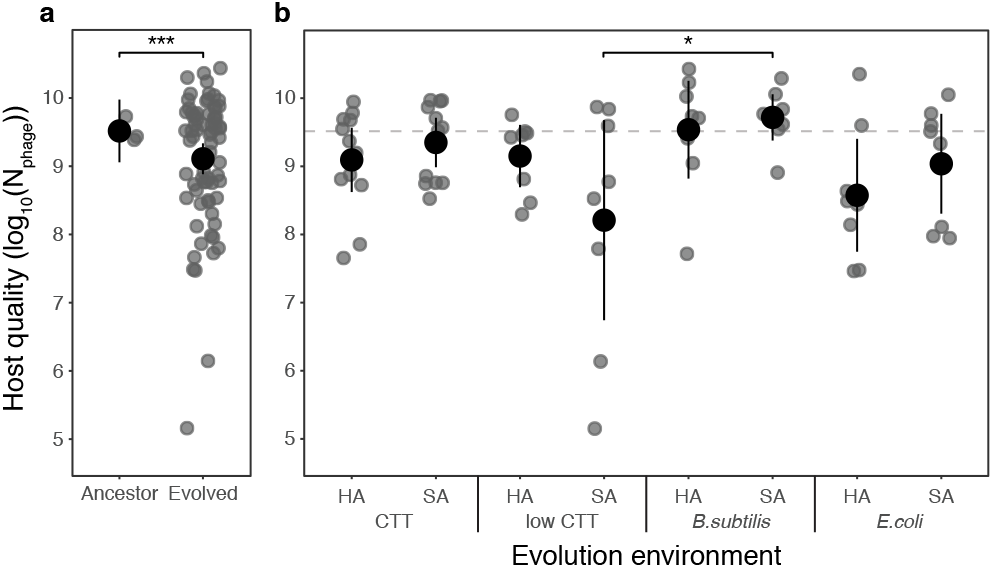
Diversification of latent host quality between independent MyxoEE-3 populations during phage-blind evolution occurred predominantly within rather than between selective environments. Cross-replicate means of host-quality measurements (grey circles) for ancestors and all evolved populations (a) and evolved populations categorized by evolution environment (b) with corresponding overall category means (black circles) and 95% confidence intervals. Host quality is measured as Mx1 phage population sizes 24 h after initial infection of bacterial populations (log-transformed data). The dashed line corresponds to average phage population size after growth on the experimental ancestors GJV1 and GJV2. The asterisks indicate statistically significant differences: two-sample two-sided *t*-test (a, *** indicates *p* < 0.001) and the one pairwise comparison in which treatment means differ significantly (b, post-hoc Tukey test, mixed linear model, * indicates *p* < 0.05)).

The observed trend of decreased host quality suggests that adaptation to laboratory conditions generally increased resistance to a major natural stress (in this case phage). This is an intriguing scenario, as laboratory “domestication” of natural isolates usually relaxes selection for natural stresses^26,75^, often resulting in corresponding trait losses^9,76,77^. Evolved bacterial populations might have become lower-quality hosts by two different mechanisms - either by being killed more rapidly by phage and thereby supporting less phage growth overall, or by individual bacteria becoming more resistant to phage. We investigate these hypotheses for one MyxoEE-3 treatment below. However, because absolute fitness is what matters from the phage perspective, our primary emphasis is on host quality *per se*, irrespective of what specific traits underlie its unselected evolution.

Despite the overall decrease in host quality across all evolved populations, no individual treatment changed significantly from the ancestor (Dunnett contrasts, all *p*-values > 0.1). This reflects variable outcomes among replicate populations within treatments (Fig 1). However, selection environment had a small but significant effect on the structure of evolved host-quality outcomes (Fig. 1b, mixed linear model, F_7,64_ = 2.4, *p* = 0.03). This effect was driven predominantly by a difference between two environments - populations that evolved with *B. subtilis* as prey on soft agar were higher-quality hosts, on average, than populations evolved on low-nutrient soft agar (Fig. 1b; post-hoc multiple comparisons with Tukey method for *p*-value adjustment t_66_ = −3.24, *p* = 0.04).

We further tested whether environmental features shared across subsets of treatments affected average host-quality evolution; we grouped treatments by nutrient type (high- and low-casitone CTT, *B. subtilis* or *E. coli*) and agar type (hard or soft agar). Agar type did not influence mean host-quality evolution (Fig. S1a, F_1,71_ = 0.028, *p* = 0.87) but nutrient type did (Fig. S1b, F_3,69_= 3.73, *p* = 0.015). The latter outcome is caused mainly by the low-nutrient and *B. subtilis* subsets, with populations growing at low nutrient levels evolving lower host quality than populations that evolved with *B. subtilis* as prey (post-hoc multiple comparison adjusted with the Tukey method, t_71_ = 2.9, *p* = 0.02).

### Degrees of within-treatment diversification varied greatly across selective environments

Although divergence among treatment means due to environmental differences was limited, we noted a high degree of diversification among populations overall that was not evenly distributed across treatments. To visualize hostquality diversification at multiple levels, we compared the coefficient of variation within vs. between selective environments for all eight treatments with A+S+ ancestors. Variation among populations within environments (on average 9.9%, ranging from 4.5 to 21.8%, Fig. 2a) greatly exceeded variation across environments (5.7%, calculated among within-treatment means, t_38_ = 4.11, *p* < 0.001), further confirming that chance differences in the mutational trajectories of replicate populations contributed more to overall diversification than did systematic differences between treatments.

**Figure 2.**
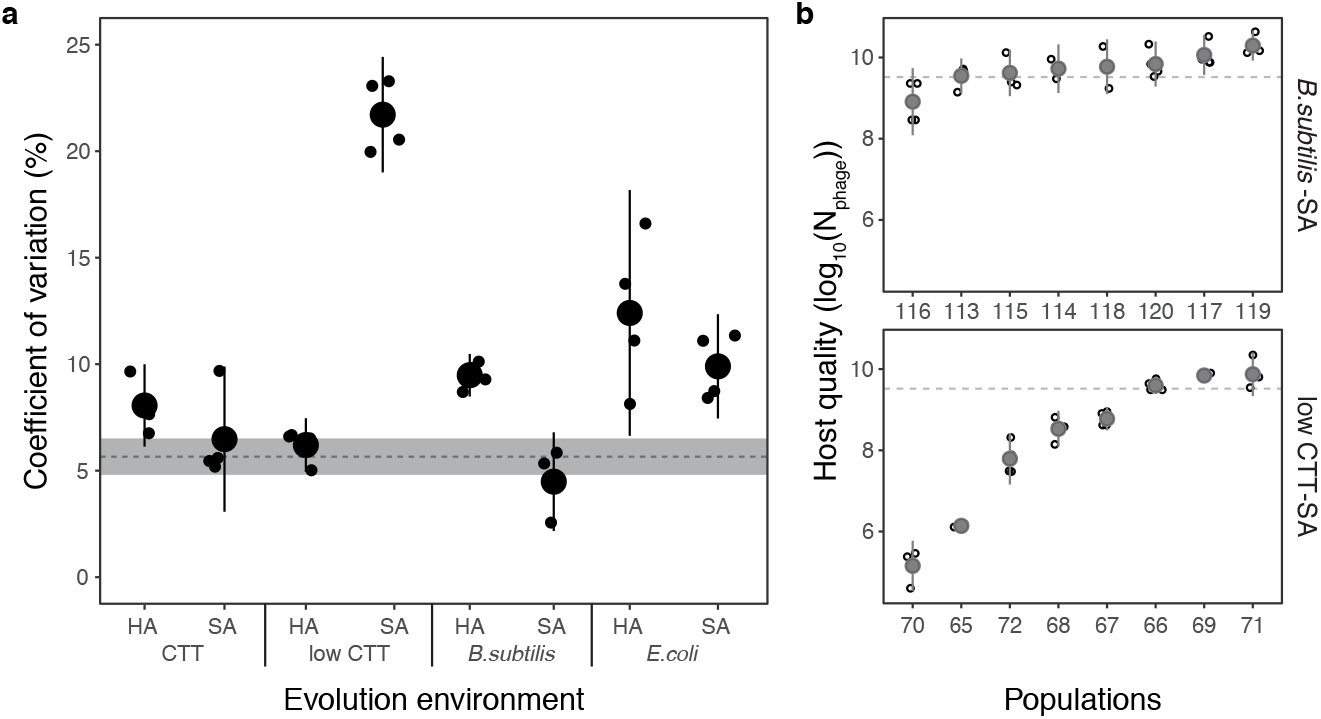
Selective environments differentially constrained stochastic diversification of host quality within treatments. (a) Within-treatment coefficients of variation (CVs) of phage population size 24 h post infection. Small and large circles represent within-replicate-assay CV estimates across evolved populations and cross-replicate-assay means for each treatment, respectively. Error bars show 95% confidence intervals. For comparison, the dashed line indicates the between-treatment CV (*i.e*. the cross-replicate-assay average of the CV among host quality mean for each treatment). The grey shaded area is the corresponding 95% confidence interval. (b) Host quality of evolved populations from the least (upper panel, *B. subtilis* prey on soft agar) and most (lower panel, low-nutrient soft agar) evolutionarily diversified treatments. Grey circles are the means across four biological replicates (open circles) and error bars represent 95% confidence intervals.

Environments might differ in the degree to which they allow latent phenotypes to diverge stochastically among replicate populations if i) LPE is caused largely by mutations that evolved due to selection rather than drift and ii) the adaptive landscapes of distinct environments differ in their ranges of accessible adaptive pathways with regard to their indirect effects on host quality. We found that the degree of host-quality diversification among replicate populations adapted to the same environment varied greatly across treatments (Fig. 2a and Fig. S2, F_7.21_ = 48.83, *p* < 0.001). At one extreme, populations evolved on soft agar with *B. subtilis* diverged very little in host quality (Fig. 2). At the other extreme, populations evolved on low-nutrient soft agar diversified much more than populations in any other treatment (Fig. 2). Such variation in diversification across environments indicates that many of the mutations underlying LPE evolved due to selection and that distinct adaptive landscapes differ in how much they constrain latent-phenotype divergence.

### Some populations latently evolved complete resistance to phage antagonism

Post-infection phage population sizes varied more than ten-fold across replicate evolved host populations within all treatments, more than 100-fold in five treatments and nearly five orders of magnitude in one treatment (low-nutrient soft agar, Figs. 1, S2). Given such diversity, we tested whether bacterial growth would be suppressed to a degree inversely correlated with phage population growth. It is not obvious that such a correlation will occur, because, as noted above, phage growth might be low on both highly susceptible and highly resistant bacterial genotypes and thereby prevent a correlation. On highly susceptible hosts, phage growth may be low because phage suppress the increase of their only growth substrate. In contrast, phage will not grow much from even large populations of highly resistant hosts. In this scenario, phage productivity could be maximal on bacterial populations exhibiting intermediate growth in the presence of phage.

During experimental epidemics with evolved populations from the most diversified evolutionary treatment (P65-P72, low-nutrient soft agar), we tracked both phage growth and *M. xanthus* population dynamics, the latter in comparison to bacterial growth in the absence of phage. None of these evolved populations grew less in the presence of phage than their ancestor, indicating that increased susceptibility to phage killing was not a general mechanism by which host quality often decreased during MyxoEE-3 (Fig. 3). As expected, these populations varied greatly in the degree to which Mx1 hindered their growth relative to their growth in the absence of phage (Fig. 3). Three evolved populations (P67, P68 and P72), like the ancestors, grew very little over the 24 h epidemics, both relative to the phage-free controls and in absolute numbers. The other five populations all grew significantly more than their ancestors, again both relative to the phage-free controls and in absolute numbers (Dunnett test against the ancestor, all *p* values < 0.005). The two evolved populations that supported the least phage growth (P65 and P70, Fig. 2b) exhibited the highest bacterial growth, which was not significantly lower than phage-free growth after 24 h. Thus, complete (or nearly complete) resistance to viral load is found to have evolved indirectly.

**Figure 3.**
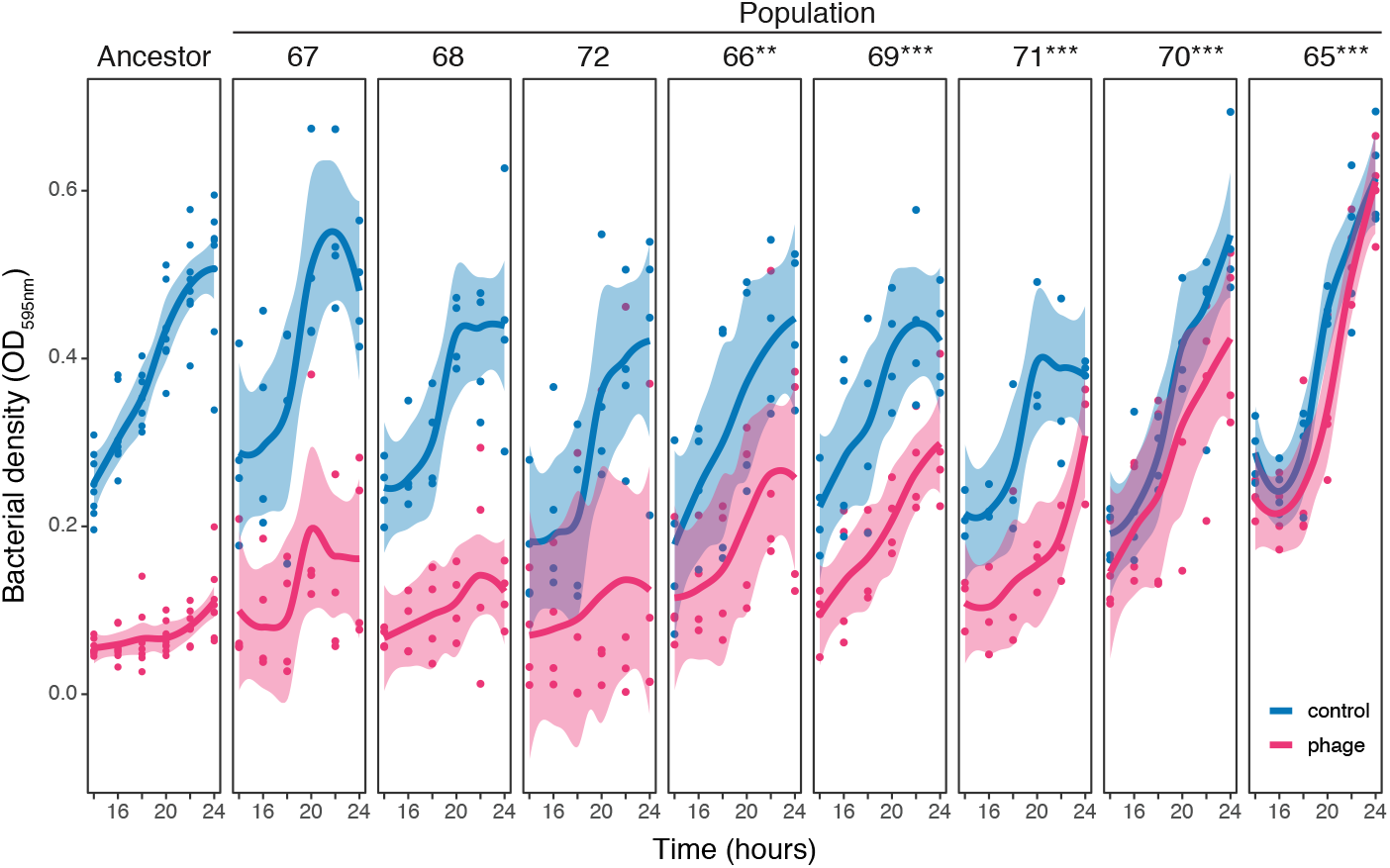
Diversity of indirect evolution of bacterial resistance to growth suppression by phage. Growth of the ancestors and of MyxoEE-3 populations evolved on low-nutrient soft agar (P65-P72) in the presence (red) and absence (blue) of phage. Data points show optical density (OD595nm) measurements over time for four temporally separate biological replicates, trendlines track conditional mean values of locally weighted regressions and grey areas represent 95% confidence intervals of the fit. The asterisks indicate significant differences to the ancestors (Dunnett test, mixed linear model, ** indicates *p* < 0.01 and *** indicates *p* < 0.001).

In contrast to the hypothetical scenario presented above, total phage productivity was found to weakly correlate with bacterial growth reduction (Spearman’s rho correlation *rs* = 0.31, *n* = 40, *p* = 0.058, Fig. 3, Fig. S3). However, large numbers of phage were able to grow from bacterial populations that exhibited very different degrees of growth suppression by phage. For example, Mx1 consistently grew to large population sizes on the ancestors, P66-P69 and P71, yet these populations varied greatly in the degree to which their growth was suppressed by phage. These results reveal idiosyncrasy in relationships between host growth and phage growth and thus point to those relationships evolving by diverse molecular mechanisms.

Intriguingly, we also noted substantial evolution and diversification of growth dynamics among the evolved populations in this treatment in the absence of phage, with several (*e.g*. P67, P68, and P71) slowing or ceasing growth earlier than the ancestors and other evolved populations (e.g. P65 and P70). Thus, variation of mutational input across replicate populations generated substantial diversification of growth dynamics in liquid media in the absence of phage.

### Ancestral motility genotype determines ancestral host quality, mean host-quality evolution, and within-treatment host-quality diversification

Motility not only allows organisms to search for new resources but also allows active flight from biotic and abiotic dangers. In bacteria, motility can allow cells to escape from non-motile phage particles^78^. On the other hand, motility-related cell-surface structures such as type-IV pili can also make bacteria susceptible to phage attack by acting as phage receptors^79^. To our knowledge, the relationship between motility and host-phage interactions, either behaviourally or evolutionarily, has yet to be examined for bacteria with multiple motility systems. We exploited the design of MyxoEE-3 - which included not only an ancestor with both *M. xanthus* motility systems intact (A+S+), but also ancestors lacking a gene essential for either motility system (A-S+, *ΔcglB* and A+S-, *ΔpilA*, Table S1) - to test for motility-genotype effects on ancestral host quality and subsequent host-quality evolution.

We quantified total phage productivity after 24 h of growth on all motility-genotype ancestors and all descendant populations that evolved on CTT hard or soft agar. The absence of *cglB* in the A-S+ ancestors had no effect on phage growth (motility effect in mixed linear model F_2,4_ = 12.47, *p* = 0.019, posthoc contrasts t4 = −0.66, *p* = 0.8), whereas the absence of *pilA* in the A+S-ancestors increased phage productivity nearly ten-fold (post hoc contrasts t_4_ = 4.79, *p* = 0.019, Fig. 4). Thus, production of pilin, but not of CglB, greatly reduces a phage epidemic.

**Figure 4.**
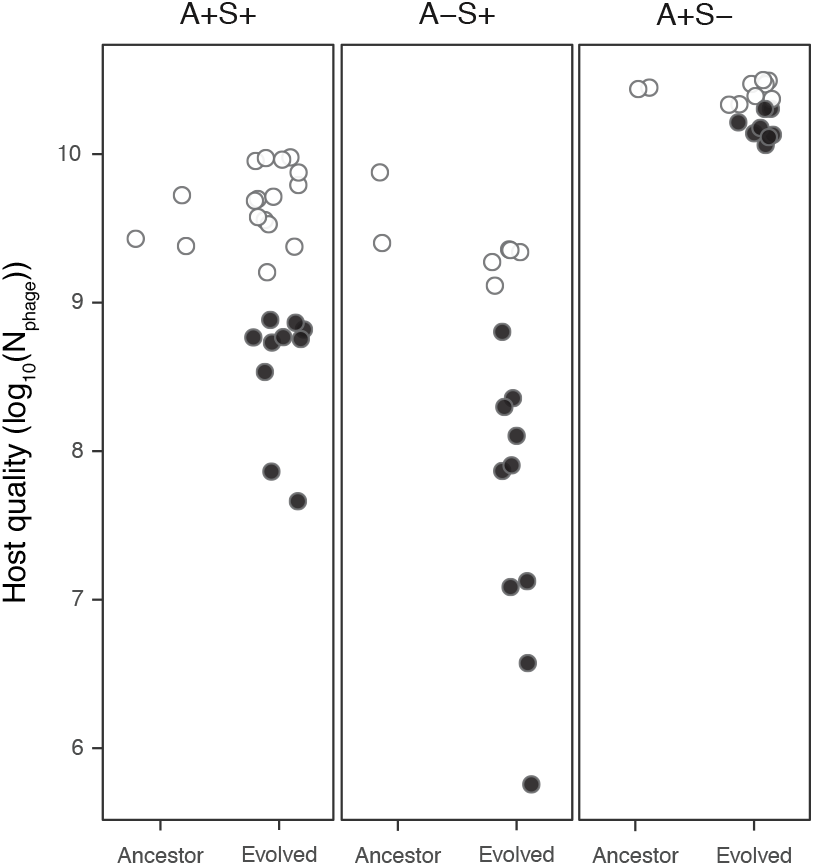
Ancestral motility genotype determines both degree of trait-mean evolution and degree of stochastic diversification for host quality. Phage population size 24 h after infection of ancestors and evolved populations with both motility systems intact (A+S+) or lacking either system (A-S+ or A+S-). Each data point represents the mean of four biological replicates. Colours show the difference between evolved populations and their respective ancestors (open circles: non-significant difference, black circles: significant difference, Dunnett test, mixed linear model).

Ancestral motility genotype also affected the character of host-quality evolution. Mean host quality of A-S+ populations decreased significantly from their ancestral values during evolution (t_55_ = 2.83, *p* = 0.007), and decreased significantly more than did the host quality of A+S+ or A+S-populations, while A+S+ populations decreased more than did A+S-populations (F_2,54_= 8.34, *p* < 0.001, all post-hoc contrasts *p* values < 0.001). Moreover, ancestral motility genotype also affected degrees of diversification within treatments, with evolved host quality spanning over four orders of magnitude among the A-S+ populations but less than a factor of ten among the A+S-populations (Fig. 4). The large differences in both ancestral phenotypes and evolutionary patterns between the A+S-category of populations vs the two categories with intact S-motility indicate that pilin production potentiates latent evolutionary reduction and diversification of host quality.

## Discussion

Previously, Meyer *et al*. found that the *E. coli* LTEE populations adapting to one selective environment in the absence of phage evolved changes in their interactions with multiple phage types^26^. Replicate LTEE populations evolved increased susceptibility toward phage T6* and increased resistance toward bacteriophage lambda with some degree of parallelism. Here we asked whether multiple distinct phage-free selective environments might differentially shape how bacterial populations descended from a common ancestor would diverge from each other in quality as phage hosts. We tested for such inter-treatment divergence with respect to both mean host quality and the degree of stochastic within-treatment diversification among replicate populations.

Overall, MyxoEE-3 populations tended to support less phage growth than their ancestors and diversified greatly, including some lineages that evolved nearly complete resistance to negative phage effects on bacterial population growth. We found high degrees of diversification in host quality among replicate populations within treatments. Indeed, among A+S+ populations, distinct selective environments drove only a small degree of divergence between selective-environment treatments in mean host quality; the primary diversifying force was chance variation in mutational input across replicate populations (Figs. 1 and 2).

Yet despite the limited effect of selection on divergence of treatment means, selection nonetheless strongly shaped the character of intrinsically non-adaptive diversification of latent phenotypes. Specifically, selective environments determined the degree to which replicate populations diversified in latent host quality, thus indicating that many of the mutations driving such divergence first evolved due to selection. Further, this result indicates that distinct adaptive landscapes can differ not only in the number of adaptive mutational pathways replicate populations might follow^60,80,81^, but can differ concomitantly in the range of latent phenotypic effects generated by those adaptive pathways^7^. Thus, distinct natural environments may often differ in the character of latent phenotypic diversity they allow to evolve.

Determination of latent stochastic diversification of host-parasite interactions by environment-specific features of fitness landscapes might apply not only to host evolution but to parasite evolution also. For example, consider a scenario with animal viruses in which different initial host species for a given viral type differ in fitness-landscape structure and thereby allow different ranges of adaptive mutational pathways to be followed by evolving viral populations. Such differences might in turn generate differences in latent virushost interaction phenotypes, including potential for jumps to novel host species (e.g. zoonosis).

We further investigated whether ancestral motility genotype is important to ancestral host-parasite interactions and/or their subsequent evolution, including extent of diversification (Fig. 4). While type-IV pili are the very means of cell invasion by phage in some host species^49,50^, we found that production of pilin, the building-block of type-IV pili^51^, both greatly reduces phage population growth in our ancestral genetic background and promotes greater latent evolutionary reduction and diversification of host quality than occurs in populations lacking pilin (Fig. 4). These immediate and evolutionary effects, respectively, of pilin production may be mediated not by pilin *per se* but by the *M. xanthus* exopolysaccharide (EPS) matrix, which is positively regulated by pilin production^69,82^. The EPS matrix is necessary for effective S-motility^83^ and mediates cell-cell adhesion^84^. We hypothesize that the EPS matrix hinders Mx1 access to its adsorption receptor (which remains unknown), thereby explaining the nearly ten-fold increase in phage growth resulting from deletion of *pilA* to generate the A+S-genotype. This hypothesis would suggest that in A+S+ and A-S+ ancestors, evolution of the EPS matrix may have often provided greater protection against phage compared to ancestral EPS, thereby promoting greater diversification than among populations lacking ancestral EPS.

It has long been recognized that forces other than direct selection on focal traits play important roles in shaping evolutionary diversification^5,13,23,85^, but latently evolved diversification is only rarely quantified^7,15,26^. We have shown that host-parasite interactions diversify greatly during parasite-blind evolution, highlighting the need to more deeply integrate LPE into our overall conception of biological diversification. The total long-term diversification of populations evolved in an original focal context can be conceived to include both divergence already actualized in that original context and the sum of all latent diversification revealed only later in new contexts. This expansive view of diversification can, in turn, inform how conservation efforts are conceived^86^ to include conservation of latent phenotypes and corresponding evolutionary potential.

## Supporting information

Supplementary Informarion

## Authors contributions

L.F. designed experiments, carried out experiments and cowrote the manuscript. M.V. designed experiments, performed statistical analysis and co-wrote the manuscript. G.J.V. designed experiments and co-wrote the manuscript.

## Acknowledgements

This work was funded in part by Swiss National Science Foundation (SNSF) grants 31003A/B_16005 to GJV and an ETH Fellowship 16-2 FEL-59 to MV. The authors thank Peter Ashcroft, Marco La Fortezza, Samay Pande, Joshua Payne, Sébastien Wielgoss and all members of the ETH Zürich Evolutionary Biology group for helpful discussions and support.

