## Supplementary Informarion for "Hidden paths to endless forms most wonderful: Parasite-blind diversification of host quality"

### Supplementary methods

*Statistical analysis*. We tested the effect of latent diversification on the final number of phage particles (log-transformed to meet normality assumptions) among evolved A+S+ populations using a mixed linear model (function lmer from the package lmerTest[1]) with the evolution environment as a fixed effect and the experimental block nested into replicates as a random factor. In addition, those populations evolved from two ancestral sub-clone genotypes of GJV1 (GJV1.1 and GJV1.2) and one sub-clone genotype of GJV2 (the rifampicin-resistant variant of GJV1). Therefore, we included the population ID nested into their ancestral background as a second random factor. However, as there was no significant difference between the rifampicin resistant and sensitive populations in final phage numbers ($F1,100 = 2.44$, $p = 0.12$), nor between the two clones of GJV1 ($t40 = 0.57$, $p = 0.57$), we combined them in the figures. To test for specific effects in the evolution environments, we used a variant of the previous model decomposing the agar type and/or nutrient types. We further measured the diversification among populations within each treatment with coefficients of variation (ratio of standard deviation to mean, multiplied by 100) and evaluated the importance of the evolution environment in shaping such divergence using the previously described mixed linear model. Bacterial growth data were analysed by i) comparing the final OD (24 h) in the presence of phage between evolved and ancestral populations (lmer model with Dunnett test, function glht from the package multicomp[2]) and ii) testing for a correlation between the phage-associated growth reduction and the phage number after 24 hours with a two-sided Spearman correlation test for paired data (function cor.test with Spearman method).

Finally, we investigated roles of both *M. xanthus* motility systems in determining host quality *per se* and its evolutionary diversification. We started by assessing the effect of motility on host quality in the ancestors using a mixed linear model with the motility system as a fixed effect and the population ID and the blocks nested into replicates as random factors. We then included the evolved populations and modified the model to include the evolutionary state (either evolved or ancestor) as a second fixed effect. We tested whether the evolved populations differed from their relative ancestors within each motility subset. All multiple comparisons were performed using the emmeans package (version 1.4.3[3]) with *p* values adjusted with the Tukey method. All statistical analyses were performed using R version 3.6.2 and RStudio version 1.2.5033[4,5]. We used the packages ggplot2[6], ggpubr (version 0.4.0) and ggsignif (version 0.6.0) to generate the figures.

**Supplementary Tables and Figures**

**Table S1. MyxoEE-3 treatments examined in this study**.

| Evolved populations* | Evolution environment | Ancestral motility genotype | Ancestral variants Rif S | Ancestral variants Rif R |
|---|---|---|---|---|
| 1 -12 | CTT (high-nutrient 1% Casitone CTT), HA (1.5% hard agar) | A+S+ | GJV1** | GJV2 |
| 13 - 20 | CTT, HA | A-S+ *ΔcglB* | GJV3 | GJV5 |
| 21 - 28 | CTT, HA | A+S- *ΔpilA* | GJV4 | GJV6 |
| 29 - 40 | CTT, SA (0.5% soft agar) | A+S+ | GJV1 | GJV2 |
| 41 - 48 | CTT, SA | A-S+ *ΔcglB* | GJV3 | GJV5 |
| 49 - 56 | CTT, SA | A+S- *ΔpilA* | GJV4 | GJV6 |
| 57 - 64 | Low CTT (low-nutrient 0.1% Casitone CTT), HA | A+S+ | GJV1 | GJV2 |
| 65 - 72 | Low CTT, SA | A+S+ | GJV1 | GJV2 |
| 89 - 96 | *E. coli* lawn grown on CTT HA | A+S+ | GJV1 | GJV2 |
| 97 - 104 | *B. subtilis* lawn grown on CTT HA | A+S+ | GJV1 | GJV2 |
| 105 - 112 | *E. coli* lawn grown on CTT SA | A+S+ | GJV1 | GJV2 |
| 113 - 120 | *B. subtilis* lawn grown on CTT SA | A+S+ | GJV1 | GJV2 |

*Odd-numbered evolved populations descend from rifampicin sensitive ancestors and even-numbered populations descend from rifampicin resistant ancestors.
**Ancestral strain GJV1 is represented by two subclones, GJV1.1 and GJV1.2 (see Methods)

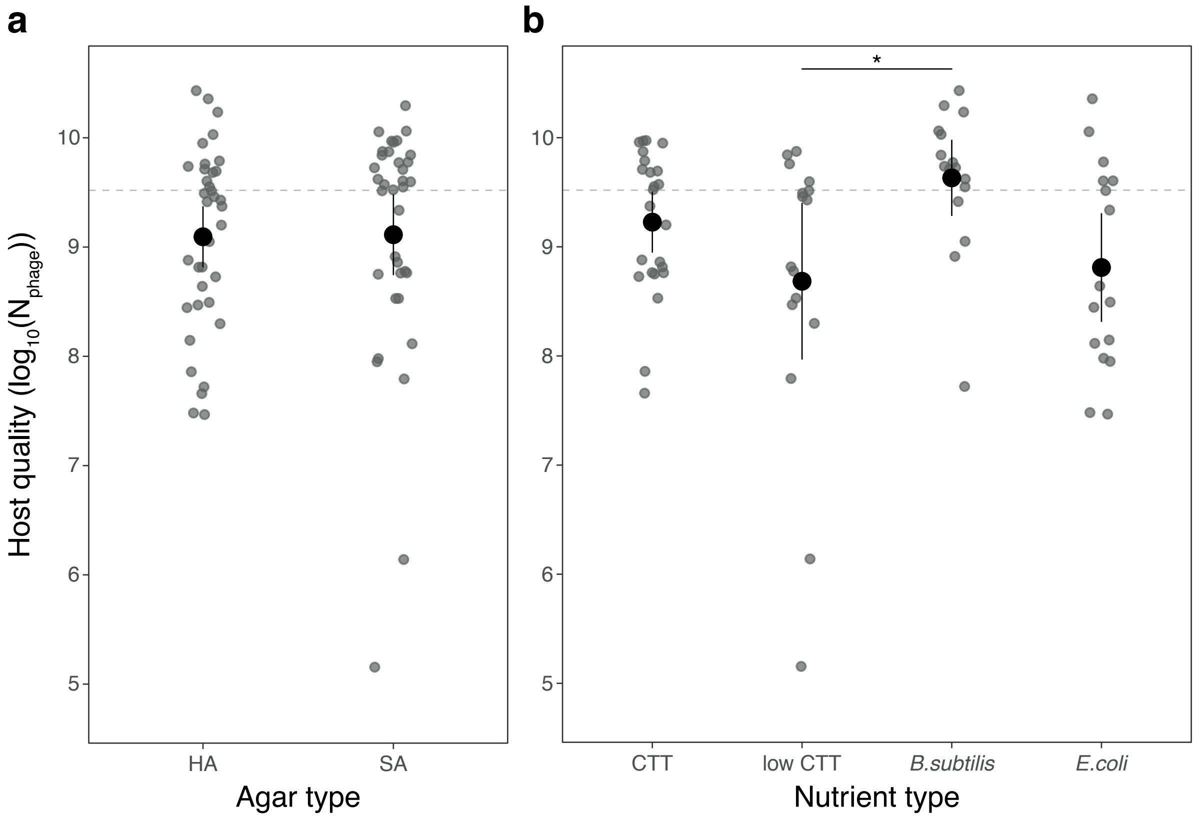


**Supplementary Figure 1. Host quality of evolved A+S+ populations grouped by shared selective-environment features.** Mx1 phage population sizes 24 h after initial infection of bacterial populations evolved on different agar (a) and nutrient (b) types. Grey circles are population means over four biological replicates. Category means (over all population means within a category) are represented by black circles and error bars are 95% confidence intervals. Dashed lines correspond to average phage population size after growth on the experimental ancestors GJV1 and GJV2. The asterisk indicates the one pairwise comparison in which category means differ significantly (post-hoc Tukey tests, mixed linear model).

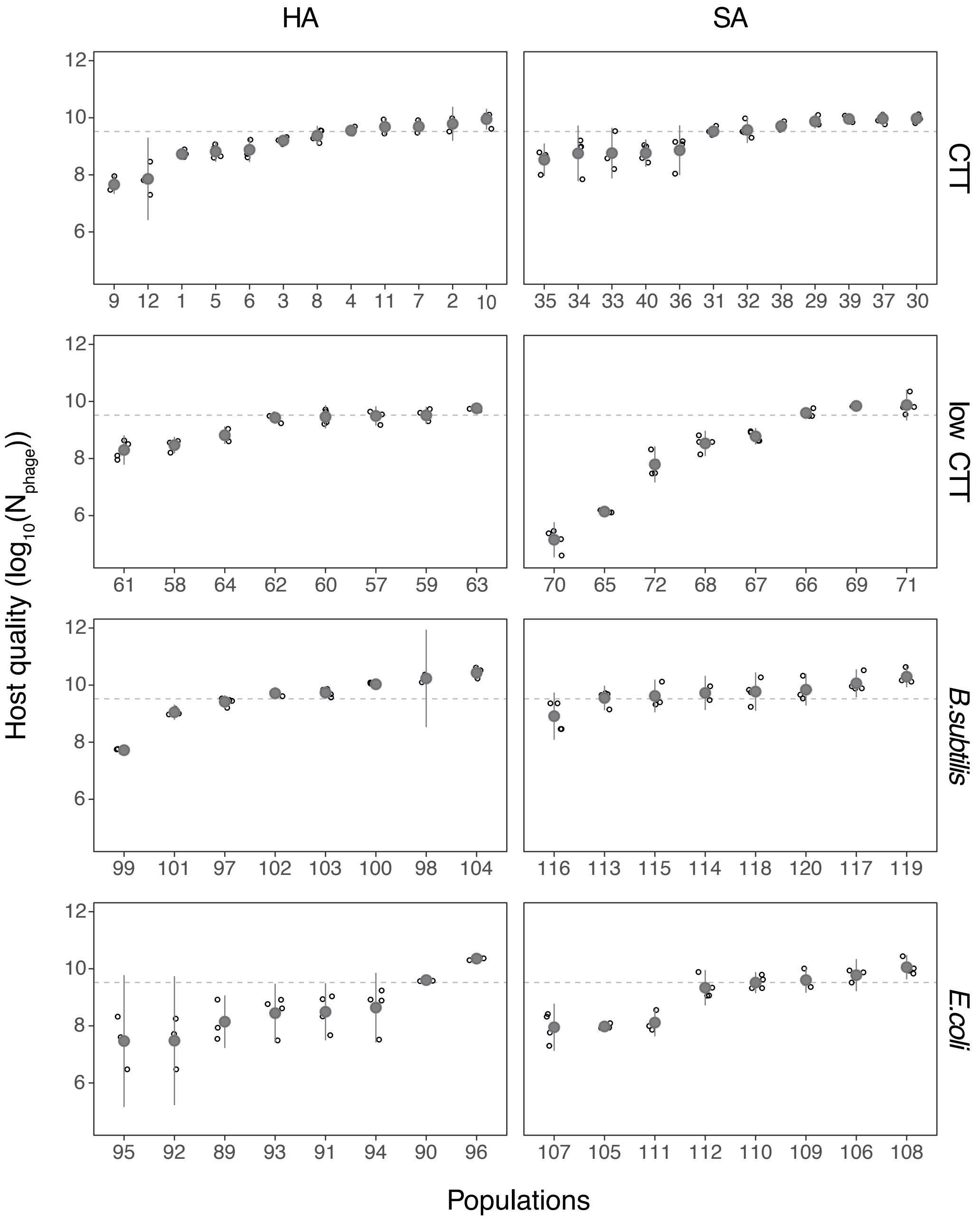


**Supplementary Figure 2. Host quality of all evolved A+S+ bacterial populations.** Host quality is measured as Mx1 phage population sizes 24 h after initial infection of bacterial populations (log-transformed data). Evolutionary treatments are categorized vertically by agar type (hard agar (HA) or soft agar (SA)) and horizontally by nutrient type (CTT, 0.1%-Casitone CTT, *B. subtilis* and *E. coli*). Open circles are original estimates per replicate, filled grey circles represent the mean, error bars represent 95% confidence intervals and dashed lines show average phage population size after growth on the ancestors.

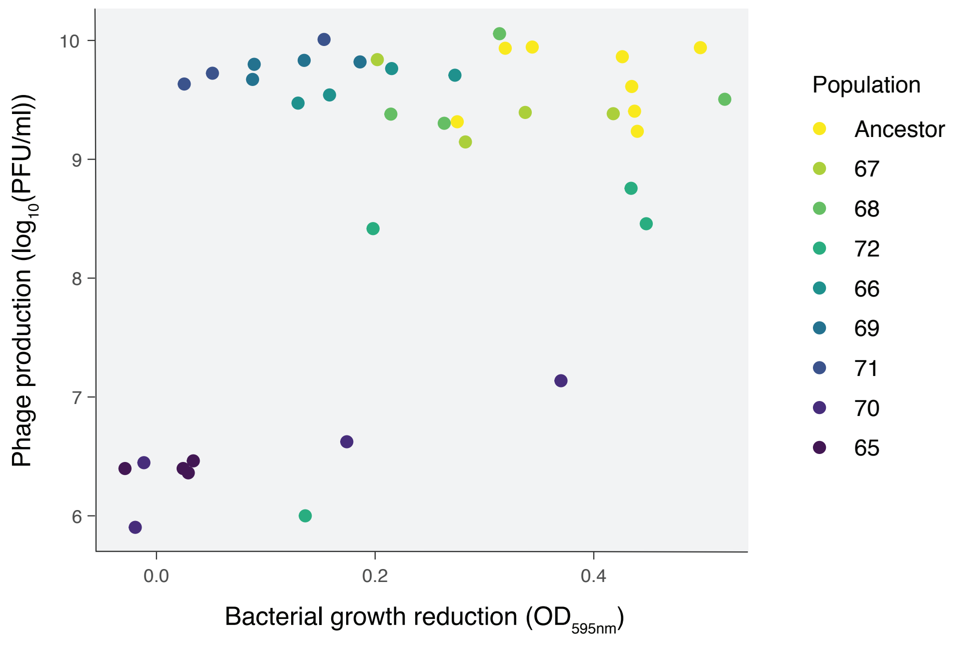


**Supplementary Figure 3. Reduction of bacterial growth by phage correlates only weakly with phage fitness.** Bacterial growth reduction is calculated as the difference in optical density ($OD_{595nm}$) in the presence and absence of phage for the ancestors and the MyxoEE-3 populations evolved on low-nutrient soft agar (P65-P72). Phage fitness is the population size 24 h after initial infection of bacterial populations (log-transformed data). Colours correspond to the different populations ($n$ = 3-8) and the colour gradient ranges from high (yellow) to low (dark purple) bacterial growth reduction.
